# Phase separation in fluids with many interacting components

**DOI:** 10.1101/2021.05.06.443002

**Authors:** Krishna Shrinivas, Michael P. Brenner

## Abstract

Fluids in natural systems, like the cytoplasm of a cell, often contain thousands of molecular species that are organized into multiple coexisting phases that enable diverse and specific functions. How interactions between numerous molecular species encode for various emergent phases is not well understood. Here we leverage approaches from random matrix theory and statistical physics to describe the emergent phase behavior of fluid mixtures with many species whose interactions are drawn randomly from an underlying distribution. Through numerical simulation and stability analyses, we show that these mixtures exhibit staged phase separation kinetics and are characterized by multiple coexisting phases at equilibrium with distinct compositions. Random-matrix theory predicts the number of existing phases at equilibrium, validated by simulations with diverse component numbers and interaction parameters. Surprisingly, this model predicts an upper bound on the number of phases, derived from dynamical considerations, that is much lower than the limit from the Gibbs phase rule, which is obtained from equilibrium thermodynamic constraints. Using a biophysically motivated model of pairwise interactions between components, we design ensembles that encode either linear or non-monotonic scaling relationships between number of components and co-existing phases, which we validate through simulation and theory. Finally, inspired by parallels in biological systems, we show that including nonequilibrium turnover of components through chemical reactions can tunably modulate the number of coexisting phases at steady-state without changing overall fluid composition. Together, our study provides a model framework that describes the emergent dynamical and steady-state phase behavior of liquid-like mixtures with many interacting constituents.

## Introduction

Fluids composed of many components with multiple co-existing phases are widespread in living and soft matter systems. A striking example occurs in the eukaryotic cell, where distinct biochemical pathways are compartmentalized into membraneless organelles called biomolecular condensates, which often form through liquid-liquid phase separation (1–3). Unlike two-phase oil-water mixtures, the cellular milieu is organized into tens of co-existing phases, each of which is enriched in specific biomolecules(1, 2, 4–8). Other prominent examples include microbial ecosystems that organize into fluid-like communities (9–11), self-assembling colloidal systems (12, 13), and synthetic multi-phase materials derived from biomolecules (14, 15). Despite their extensive prevalence, our understanding of how microscopic interaction networks between individual constituents encode emergent multi-phase behavior remains limited.

Delineating the co-existing phases of a heterogeneous mixture is a problem with a rich history (16) – determined by constraints of chemical, mechanical, and thermal equilibrium. In mixtures with few components (fewer than 5), a combination of theory, simulation, and experiment has enabled extensive characterization of phase separation kinetics and equilibrium co-existence (17–23) and the interplay between phase separation and chemical reactions (19, 24, 25). In the biological context, recent studies have begun to connect biomolecular features to their macroscopic phase behavior in binary or ternary mixtures (7, 26, 27). However, as the number of components increase, determining the emergent phase behavior from the underlying constraints becomes unwieldy and intractable – from both analytical and numerical standpoints except for very particular systems such as polydisperse blends of a single species (28). An alternate approach, originally proposed by Sear and Cuesta (29), aims to characterize the phase behavior of mixtures that contain many components whose pairwise interactions are drawn from a random distribution. By building on results on properties of random matrices, originally identified by Wigner (30) and subsequently applied in various contexts (31–33), they relate the initial direction of phase separation to properties of the interaction distribution, subsequently confirmed independently by simulation (34). These results, however, are limited to describing only the *initial* direction of phase-separation for *marginally* stable fluid mixtures i.e. coinciding exactly at the spinodal. Consequently, little is known about the overall phase behavior of fluid mixtures that spontaneously demix i.e. within the spinodal – including kinetics beyond the initial direction of phase separation or the number and composition of coexisting phases at equilibrium. More generally, the emergent phase behavior of fluid mixtures with many randomly interacting components is not well understood. This lack of understanding, in turn, limits our ability to rationally program fluid mixtures with different macroscopic properties.

Here, we develop a dynamic model of phase separation in fluid mixtures with many randomly interacting components. Through simulation of the model, we demonstrate that fluid mixtures with many components exhibit characteristic similarities in phase-separation kinetics and in the number and compositional features of co-existing phases at steady-state, even when the underlying interactions are random. We propose a simple model, combining insights from random-matrix theory and dynamical systems analyses, that predicts dynamical and steady-state characteristics of the emergent phase behavior. By constructing a biophysically motivated model of the pairwise interaction distribution, we discuss two distinct ensembles (or component design strategies) that encode either linear or non-monotic scaling (i.e. with an optima) between the number of co-existing phases and components. Finally, we extend our framework to incorporate chemical reactions, and show that active turnover of components can tunably modulate the number of co-existing phases at steady state even without altering overall fluid composition. Overall, our model provides a framework to predict and design emergent multi-phase kinetics, compositions, and steady-state properties in fluid mixtures with many interacting components.

## Results

### Model definition

We begin by describing the free-energy of a mixture of (*N* + 1) interacting species through a mean-field regular solution model at fixed temperature and overall volume (Eq. 1). Here, *ϕ_i_* represents the volumefraction of each species *i*, and *ϕ_s_* = 1 – ∑_*i*_*ϕ_i_* is the volume fraction of the remaining component, which is typically the solvent. Equivalently, this model can also represent the *normalized* volume fractions of (*N* + 1) components in a solution where the total solute concentration is invariant across phases. *χ_ij_* and *χ_is_* are the effective pairwise interactions amongst different species and between components and solvent respectively. For simplicity, we assume an initially uniform solute mixture (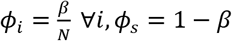; *β* is total solute volume-fraction), inert solvent (*χ_is_* ≈ 0) and components that don’t self-interact i.e. *χ_ij_* only depends on interaction-energy (*ϵ_ij_*) between components *i, j, i* ≠ *j.* Further, we stipulate that the pairwise interactions *χ_ij_* are independent random variables which are drawn from a distribution with finite mean *v* and variance *σ*^2^ (Figure 1A).

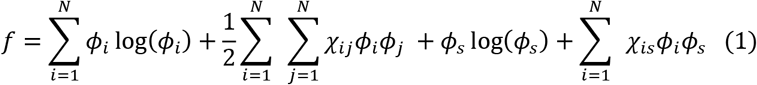

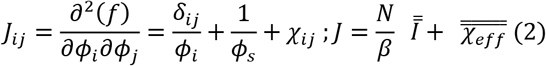

**Figure 1.**
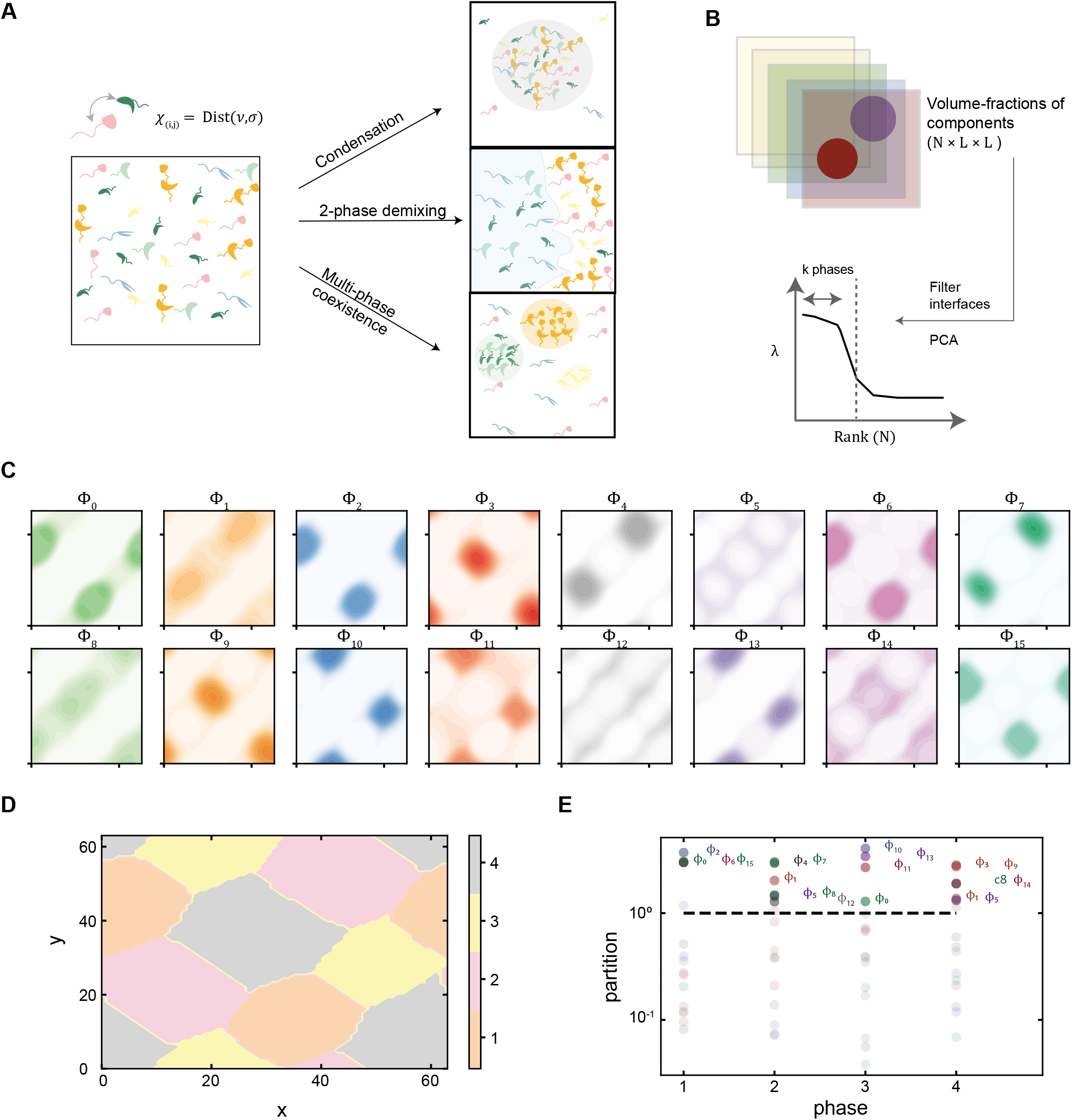
A model for phase separation in multi-component fluid mixtures. **(A)** A schematic depicting that the interactions between pairs of components are randomly drawn from a distribution and encode varying emergent properties. **(B)** Schematic depicting post-processing analyses on simulation data to identify the co-existing phases from PCA **(C).** Plots depict volume-fraction profiles of 16 components (labeled *c*0 to *c*15) at steady-state from a single trajectory with identical color-bar scales (0,0.75). Darker colors represent regions of higher volume-fraction and simulation parameters are presented in main text and SI. **(D).** The different phases (labeled 1 till 4) present at steady-state in (A) are depicted here. **(E).** The partition-ratio (ratio of average volume-fraction in a phase over total initial volume-fraction) of all components are plotted for each phase (x-axis) at steady-state conditions shown in (A). The highlighted components are enriched in those respective phases and the dashed-lines represent no enrichment (partition=1).

The point beyond which a mixture spontaneously phase-separates - the spinodal, or the *marginally* stable state -- occurs when the minimum eigenvalue *λ_min_* of the Jacobian matrix *J* (Eq. 2; SI Appendix) crosses zero i.e. *λ_min_*(*J*) = 0. The corresponding eigenvector gives the initial direction of instability (Fig S1A-C, SI Appendix), which either leads to condensation-type (*v* ≤ −*N*) or demixing-type instabilities (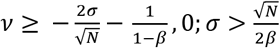) depending on the values of *v,σ* (29, 34). During condensation, the instability points toward dilute and dense phases with similar compositions since individual components are strongly attractive on average, as determined from the angle between the marginal eigen-vector and the initial composition being close to 0 *or* 180 (Fig S1C). Conversely, during de-mixing, the initial instability, whose direction is roughly perpendicular to the uniform mixture (Fig S1C), points to phases with distinct compositions. In general, solutions that de-mix contain unstable modes beyond the marginally stable point, potentially leading to multi-phase co-existence. Multiphase coexistence does not generically occur in condensation transitions, because of the band gap between the smallest eigenvalue and the rest of the spectra (Fig S1, SI Appendix). Motivated by this, here we focus on phase separation in solutions whose component-interactions are variable, but not strongly attractive on average (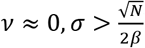).

After the initial instability, fluid mixtures undergoing spinodal decomposition display rich dynamics and diverse multiphase co-existence (Figure 1A). To probe the kinetics and emergent equilibrium properties, we formulate a set of dynamical equations to track the evolution of *N* independent volume fractions 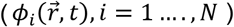 (Eq. 3). The temporal evolution of a component’s volume-fraction 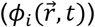 depends on diffusive fluxes driven by gradients of chemical potential 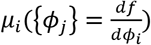 with a mobility coefficient *M_i_*, = *M_ϕ_i__* also known as conserved model B dynamics (35, 36). We assume that all components obey *M_i_* = *M_ϕ_i__*, approximately recapitulating Fickian diffusion in the limit of non-interacting, dilute components (SI Appendix). Finally, we include surface-tension effects ensuring long-wave-length stability by modifying the bulk chemical potentials with a component-independent gradient term 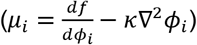.

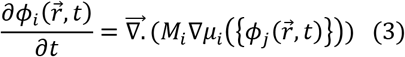

We numerically simulate these non-linear, coupled partial differential equations using Fourier space representations to compute gradients and fluxes (SI Appendix). Unless specified otherwise, a 2D grid (*L* × *L, L* = 64) is initialized with a uniform and equimolar solution 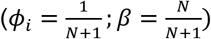 with small compositional fluctuations. For each simulation, we sample the interaction matrix *χ* from a normal distribution (Figure 1A) with zero mean and specified variance (quenched disorder). At any time point, the state of the system is described by the volume-fraction matrix (*N* × *L* × *L*), giving the volume fraction of each component at every point in space. We can infer the number of phases *N_ph_* by performing PCA on this matrix after filtering out interfaces between phases that vary in composition. We then identify the significant eigen-values which correspond to individual phases (Fig 1B, SI Appendix). We identify which phase each point in space belongs to by first performing K-means clustering (with the number of clusters as the number of phases from PCA) followed by classification to assign individual points to the closest phase by composition 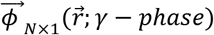. For all the points assigned to a phase, we compute spatially-averaged volume-fractions that characterizes the bulk composition of the *γ* phase 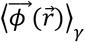 (SI Appendix).

### Multiple phases with distinct compositions characterize random fluid mixtures

To explore phase-behavior of multi-component solutions, we first simulate the dynamics of an initially equimolar solution of *N* = 16 components (labeled *ϕ_i_*) whose pairwise interactions are randomly sampled from *χ_ij_* ~ Normal(0, *σ* = 4.8). This choice of parameters ensures spontaneous phase separation with multiple initial unstable modes. At steady-state, the mixture exhibits heterogeneous, multi-phase coexistence (Figure 1C). The steady state solution has 4 co-existing phases (Figure 1D) that are enriched in a distinct yet characteristic number of components per phase (Figure 1E; SI Appendix). Such multi-phase coexistence with differing compositions occurs generically for different choices of parameters (Figure S2).

A key question is to understand how the steady state properties of the phases relate to the interaction distribution between components. To explore this, we ran many simulations under identical conditions while re-sampling from the interaction matrix specified above. Although the precise values of steady state compositions vary between different simulations, there were striking statistical similarities between both the number of distinct phases at steady state, and the temporal dynamics leading to this steady state (Figure 2A; green line is trajectory in Figures 1C-E). This statistical convergence in the expected number of co-existing phases is independent of the choice of mobility parameter (Figure S3B), simulation length (Figure S3C), or specific simulation parameters (Figure S3D).

**Figure 2.**
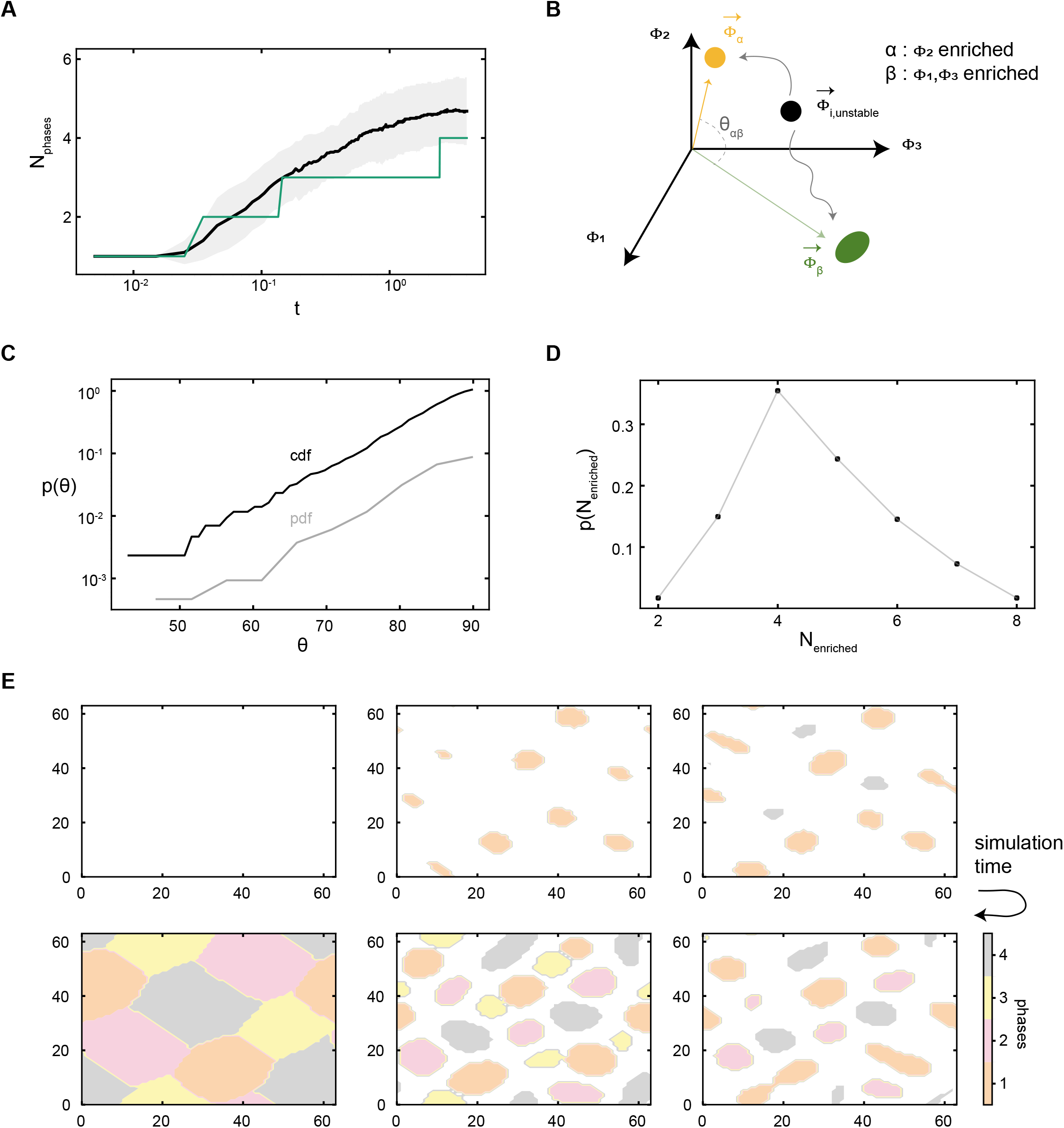
Multiple phases with distinct compositions characterize random fluid mixtures. **(A).** Number of co-existing phase (y-axis) versus simulation time (x-axis, log-scale) for simulation conditions as in Fig 1C. The solid line represents mean of 50 different trajectories, the filled regions represent 1 standard deviation, and the green line represents the specific trajectory whose steady-state properties are shown in Figs 1C-E. **(B).** Schematic illustrating how compositional observables are computed from steady-state compositions i.e. angle between co-existing phases (*θ*) and number of enriched components per phase are computed. In the example, an initially unstable phase of 3-components de-mixes to form two-phases (*α,β*) that are enriched in distinct number of components (shown in legend). **(C).** Probability (pdf) and cumulative distribution (cdf) of angles between co-existing phases at steadystate for simulation parameters in (D). **(D).** Probability (*p_N_enr__*) distribution of number of enriched components (x-axis, *N_enr_*) per phase at steadystate for simulation parameters in (D). **(E).** Individual snapshots of the simulation trajectory reported in Fig 1C labeled with existing phases (1-4 from steady-state, white or unlabeled if like initial equimolar solution). The steady-state labels are shown in the colorbar whose bulk compositions are used to assign phase labels at earlier times (SI Appendix).

The compositions of the steady state phases also exhibit similarities. To characterize the compositions at steady-state, we compute the angle (Figure 2B) between compositions of the pairs of co-existing phases (*θ_αβ_*) and also measure the number of components enriched in each particular phase (*N_enriched_*). Phases with *θ_αβ_* close to 0 have largely similar compositions, while those close to π/2 are enriched in different sets of components. Figure 2C shows the distribution *p*(*θ_αβ_*) calculated across multiple simulations, demonstrating that different phases have composition vectors that are orthogonal. Since individual concentrations must be positive, this indicates that co-existing phases are enriched in distinct sets of components. The number of enriched components per phase *p*(*N_enriched_*) is distributed around values of *N_enriched_* = 3,4,5 (Figure 2D). This is consistent with the distinct phases being orthogonal in component partitioning 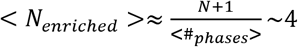. This observed compositional orthogonality is independent of the specific simulation parameters (Figure S3E-F). Overall, the equilibrium phase behavior of random fluid mixtures is characterized by multiple co-existing phases with distinct compositions.

### A simple random-matrix derived theory predicts equilibrium phase behavior of fluid mixtures

The consistency in dynamic and equilibrium properties of random mixtures motivated us to explore whether we could unify the emergent phase behavior through a theoretical framework. First, we asked whether the equilibrium multi-phase co-existence was related to the properties of the initially uniform mixture, as characterized by the Jacobian matrix. We ran simulations across several conditions (varying *σ,N*), and computed both the number of unstable modes at the *beginning* of each trajectory (*N*_*λ*<0_ number of negative eigen-values from linear stability analyses, SI Appendix) as well as the number of co-existing phases at *equilibrium* (*N_ph_*). Strikingly, when examined across simulations with diverse parameters, these exhibited a linear relation where *N_ph_* ≈ *N*_*λ*<0_ + 1 (Figure 3A). This implies that each linearized unstable eigenmode typically gives rise to a *unique* co-existing phase at equilibrium (SI Appendix). Since the eigen-vectors corresponding to these initial unstable modes are largely perpendicular to each other (Figure S4A; SI Appendix), this may contribute to the observed compositional orthogonality between coexisting at steady-state (Figure 2C, Figure S3E). Overall, our results suggest that the equilibrium number of co-existing phases in fluid mixtures undergoing spontaneous phase separation can be computed by simply computing the number of unstable modes in the uniform mixture. This is a striking conclusion because in general the number of stable equilibria in a nonlinear free energy functional is independent of the number of unstable modes in the initial dynamics.

**Figure 3.**
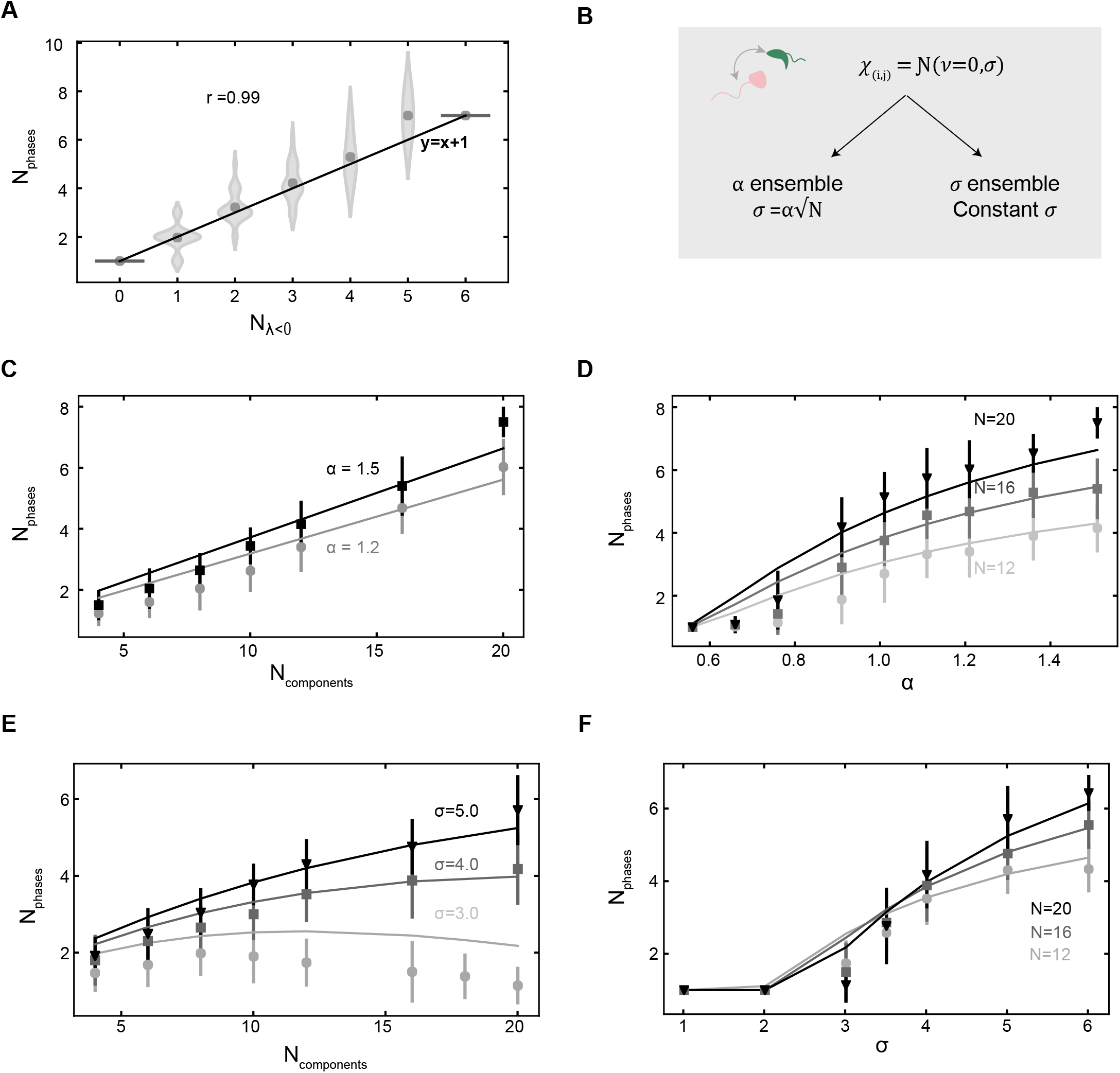
A simple scaling predicts equilibrium behavior of random mixtures from different ensembles. **(A).** Number of phases at steady-state (y-axis) versus number of negative eigen-values of the initial equimolar mixture. Histograms represent simulation results that are collapsed from a range of different *N, σ* and solid line represent the equation *y* = *x* + 1. The correlation coefficient is reported between the mean of simulation results and the solid line. **(B).** Schematic depicting the two interaction ensembles with different variance scaling. The *a* ensemble consists of components whose variance in interactions scales with number of distinct species and the *σ* ensemble has a distribution of fixed variance. A biophysical model to motivate these two ensembles is discussed in text and illustrated in Figure S4A. **(C-D).** Variation of number of co-existing phases at steady-state with number of components (linear-scaling) and different values of *α* (monotonic saturation) in the *α* ensemble. Solid lines represent theoretical predictions, dots represent mean of simulation results, and vertical dashes represent one standard-deviation around the mean. In each plot, darker lines represent higher values of *α* and *N* respectively, and different marker types are employed to reinforce this. **(E-F).** Variation of number of co-existing phases at steady-state with number of components (non-monotonic) and different values of *σ* (monotonic saturation) in the fixed *σ* ensemble. Solid lines represent theoretical predictions, dots represent mean of simulation results, and vertical dashes represent one standard-deviation around the mean. In each plot, darker lines represent higher values of *σ* and *N* respectively, and different marker types are employed to re-inforce this.

The number of unstable modes or negative eigenvalues can be counted using Wigner’s semi-circle law for random matrices, giving 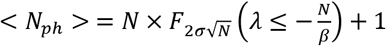 (SI Appendix). Here, *F* is the cdf of the semi-circle distribution whose eigen-values are between 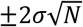 and the argument is the entropic cost that needs to be offset to phase separate (SI Appendix). Since the eigenvalues of the semi-circle distribution are equally spaced on average, the number of phases can be approximated as

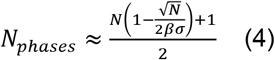

Eq. (4) implies that when 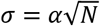, the number of phases scale linearly with the number of components. In the next section, we will discuss a biophysical interpretation of the proportionality constant *α*. When 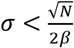, there are no unstable modes and the uniform phase remains stable, as expected. If the interactions between species are strongly variable 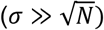, Eq. (4) implies a maximum of 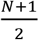 coexisting phases at equilibrium. Interestingly, this asymptotic scaling is significantly less than the upper constraint of *N* + 2 (or *N* if temperature and pressure/volume are fixed) originally formulated by Gibbs (16). Note that this asymptotic scaling of *N*/2 will continue to hold even when the average of the interaction distribution, *v,* is non-zero, which at most adds only one single orthogonal eigen-mode (SI Appendix). After the initial instability, linear stability analyses predicts that each unstable mode grows exponentially (exp -*αλt*) (SI Appendix), so the characteristic time for a phase to form scales as *t_ph_* ∝ 1/*Λ*. Since the unstable eigen-values are equally spaced on average (*λ_min_* = < *λ*_2_ < ⋯ < *λ_k_* < ⋯ < *λ_γ_* < 0 *s. t. λ_k_* – *λ*_*k*+1_ = *constant*), the typical time for the *k^th^* phase to macroscopically form is larger for higher *k* is 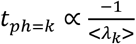 (the proportionality constant can be approximately estimated, see SI Appendix). This relation, though an approximation, predicts multi-staged phase separation kinetics: newer phases macroscopically emerge at later times in a sequential order – predictions that are consistent with observations (Figure 2A, Figure 3D, Figure S4B). This is made vivid by tracking the temporal evolution of different phases for the example trajectory shown in Figure 1C (Figure 2E; SI Appendix). Together, these results support a model derived from random-matrix theory that connects statistical properties of the initially homogeneous solution to the dynamics, compositional features, and number of steady-state phases in fluid mixtures with randomly interacting components.

### A biophysically motivated model for component interactions predicts ensembles that encode linear or optimal scaling of co-existing phases with number of components

We define a simple biophysically motivated model of interactions (Figure S5A) to explore encoding of multi-phase behavior. In this model, each component has *L* interaction sites that each exhibit one of two characteristics (*A or B*), for example negative or positive charges or binding and receptor domains, with favorable interactions between unlike sites and unfavorable interactions between like sites. This might reflect an ensemble of multivalent proteins with different interacting domains, a mixture of polypeptides with different charge sequences, or DNA-coated colloidal systems with varying sequence features. We consider a component mixture where each type of site is equally probable, and the attractive and repulsive interactions have the same energy-scale *ϵ*. The average interaction between two randomly sampled species is then *v* = ∑_*L*_ < *ϵ_ij_* >= 0, and the variance of interactions is 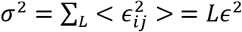. Hence, the distribution of interactions between two random components is binomial, and well-approximated as gaussian with 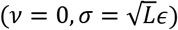. This argument implies two different ensembles (Figure 3B): one in which mixtures have a fixed number of sites (*L*) and thus fixed *σ* (the *σ* ensemble), and another (the *α* ensemble) where the number of interacting sites scale with number of components 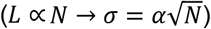. These ensembles should exhibit distinct encoding relationships between number of coexisting phases and number of components.

To explore this, we ran simulations across a wide range of (*N,α,σ*). In the *α* ensemble (Figure 3B), the predicted scaling of number of co-existing phases from Eq (3) scales linearly with *N*, and saturates with increasing *α*, in agreement with theoretical predictions (Figures 3C-D). By contrast, in the *σ* ensemble, theory predicts an optimal number of components that maximizes the number of co-existing phases at equilibrium 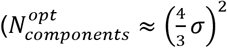; SI Appendix), which broadly agrees with simulation predictions (Figure 3E-F). Intuitively, with fewer components, phase separation is promoted due to lower entropic costs, but the maximum number of co-existing phases is bounded by 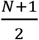. Conversely, in the limit of many components, the system is stable and doesn’t phase separate (entropic stabilization ∝ *N* whereas enthalpic terms scale as 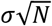), thus leading to non-monotonic scaling. In all cases, the predictions deviate from simulation at either low *N* or *σ, α*, when fluctuations in the unstable modes of the eigen-spectra are of order unity. Finally, simulations match theory for equimolar solutes with lower total solute volume-fractions (lower *β,* Figure S5B) and theoretical predictions continue to exhibit similar scaling relationships (Figures S5C-F) in regimes where simulations are numerically inaccessible 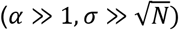. These results show that increasing the variance of interactions proportionally with the number of components, for example by increasing the number of interactions sites, allows encoding more phases. Conversely, sampling more components from a distribution of fixed variance, for example all components have the same number of interaction sites, has a maximal number of co-existing phases.

### Active turnover of components modulates number of co-existing phases at steady state

In most biological systems, interacting components are actively being produced and degraded (37). To study how such chemical reactions impact phase behavior, we modify our dynamical equations to:

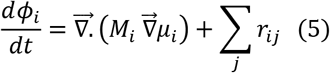

where *r_ij_* are set of reactions ({*j*}) that change the fluxes of species *i.* In the simplest case of turnover, each component is produced at fixed rate *k* and degraded at rate *k_off_* = *kN*/*β*. In the absence of phase separation, each component’s steady-state volume-fraction is 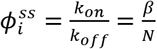. As before, when 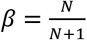, this corresponds to an equimolar solution i.e. 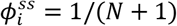.

We then perform linear stability analyses (SI Appendix) which shows that phase separation is suppressed by high rates of turnover i.e. larger values of *k_off_*, consistent with previous theoretical work in binary or ternary mixtures (19, 24). Active turnover effectively introduces local mixing by cyclically synthesizing and degrading components which counteracts spatial variations that arise from phase-separation. When the rate of turnover becomes dominant to induce mixing at large length-scales, it effectively decreases the band of unstable eigen-values that contribute to phase separation. The unstable eigen-values that continue to persist are orthogonal to each other and drive multi-phase co-existence albeit with fewer number of steady-state phases. More generally, the higher the rate of turnover, the fewer number of steady-state phases (SI Appendix). Further, our theory predicts that the number of co-existing phases at steady-state can be tunably suppressed by varying the absolute rates of turnover (*k_on_, k_off_*), even when keeping their ratio constant i.e. the overall fluid composition at steady-state remains equimolar and identical to the initial conditions, with a scaling of 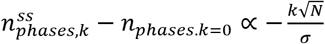 (SI Appendix). To test this hypothesis, we ran simulations in which we varied the rate of turnover while keeping their ratio constant (increasing both *k_on_, k_off_* for all components), and averaged observables across replicate trajectories. We find that increasing rates of active turnover leads to decreasing number of co-existing phases (Figure 4A) and simulations largely agree with the simple theoretical prediction (Figure 4B). This suggests that active turnover of components can serve as a route to tunably modulate multi-phase co-existence in random fluids even without altering the relative or overall composition of such mixtures.

**Figure 4.**
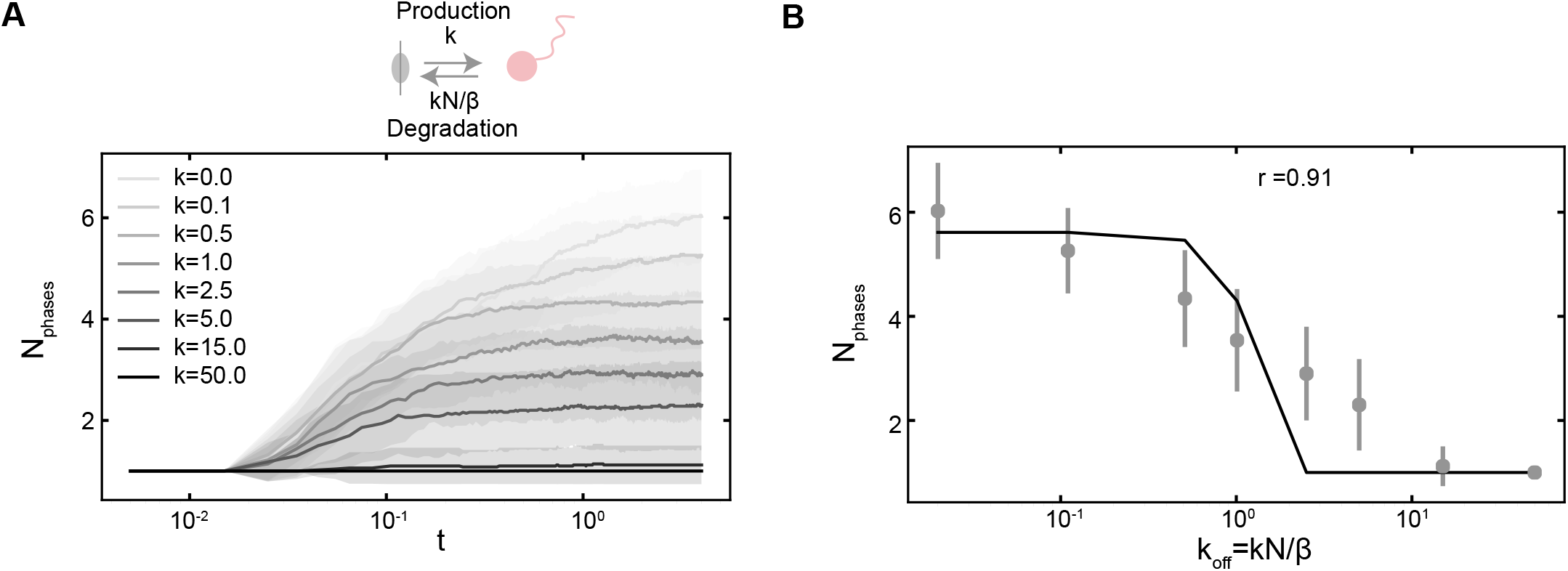
Active species turnover tunably modulates multi-phase coexistence at steady-state. **(A).** Schematic depicts constant production and first-order degradation of components. The graph shows number of phases vs simulation time (in log-scale) across a range of reaction rates in a system with *N* = 20 components and *σ* = 5.4. Solid lines represent mean of trajectories (>40 replicates per condition), shaded bars represent 1 standard deviation on each side of the mean, and darker lines corresponds to faster rates of turnover. **(B).** Simulation (dots) and theoretical predictions (solid line) on number of co-existing phases at steadystate versus rate of turnover. *r* value indicates correlation coefficient between theory and simulation, solid-circles represent mean number of coexisting phases, and vertical lines represent one-standard deviation ranges under the same parameter conditions as above.

## Discussion

Over the past several years, there has been a growing appreciation of the role of multicomponent and co-existing phases inside a cell. These phases, or condensates, compartmentalize many interacting species and pathways to enable diverse yet specific functions across cell-types and organisms. More generally, fluid mixtures with many phases and components are prevalent in biology, soft-matter, and industry. Yet, we still do not understand how numerous interacting components encode the emergent multi-phase behavior. The goal of this study is to develop a simple model of the dynamics and equilibrium phase behavior in fluid mixtures with many components. We choose the interactions between components from an underlying distribution, and thereby can use Random Matrix Theory to analyze the resulting dynamics. Through simulation and theory, we find that spontaneous phase separation of such mixtures is characterized by staged phase separation dynamics, and multiple co-existing phases at equilibrium with distinct non-overlapping compositions. Importantly, our model suggests that these characteristics do not require fine-tuning of composition or interaction parameters, rather, they are an emergent property of fluid mixtures with many components with random interactions. By formulating a biophysically motivated model of pairwise interactions, we design different component ensembles that encode linear or optimal scaling of number of co-existing phases versus components, which we validate through simulation and random matrix theory. Strikingly, we identify an upper bound for the maximum number of co-existing phases in random mixtures, derived from dynamical considerations, that is asymptotically lower than the Gibbs phase rule. Random interactions effectively introduce competing interaction networks, analogous to the concept of “frustration”, which likely limits the maximal number of possible co-existing phases. Motivated by the observation that biological fluids often exhibit component turnover such as synthesis/degradation of biomolecules, we show that active turnover of components can tunably modify steady-state multiphase co-existence, even without altering overall fluid composition.

The model formulated herein is only the first step in being able to *design* multicomponent phases in terms of their individual components. Recently, there has been tremendous progress in characterizing the sequence to phase-behavior relationship of individual proteins and nucleic acids (14, 26, 38), composition of different condensates (39–41), and regulated formation of condensates at specific locations, often through nucleation (42, 43). Soft-matter colloidal systems (44, 45), DNA-based nanotechnology (12, 46), programmable magnetic materials (47), and multiplexed protein-design offer diverse attractive routes to both experimentally test predictions and serve as platforms to enable design of multi-phase fluid mixtures. Leveraging random-matrix theory approaches, as done in our study for spontaneous phase separation and in a recent related pre-print studying nucleation in metastable fluids (48), will enable programmable design of multi-phase co-existence in multicomponent fluid mixtures. An important related problem is the design of targeted multi-phase mixtures whose compositions and interactions are specifically tuned, not random. Computational and theoretical approaches for programming phase behavior in these systems will enable material design using synthetic and biological constituents. Another exciting direction is to incorporate energy consuming processes as part of the design. Examples include non-reciprocal interactions, chemical reaction networks, molecular motors, and motile particles – all of which are characteristic of living systems. More generally, studying the interplay of non-equilibrium processes and multi-phase behavior in fluid mixtures will be an exciting and rich area for biology and soft-matter physics.

## Materials and SI Appendix

Phase-field simulations and subsequent data-analyses were performed using custom code written in *python.* We employ results from random matrix theory and dynamical systems analyses in deriving the theoretical scaling relationships presented in the text. More details about theory, simulation, and numerical methods for post-processing are available in the Supplementary Information.

## Data availability

All the code used to run simulations will be available on a publicly accessible github repository.

## Acknowledgments

We thank Ofer Kimchi, Ella King, Jon Henninger, and members of the Brenner lab for helpful discussions. K.S was supported by the NSF-Simons Center for Mathematical and Statistical Analysis of Biology at Harvard (award number #1764269) and the Harvard FAS Quantitative Biology Initiative. MPB was supported by ONR N00014-17-1-3029 and the Simons Foundation.

