## Supplementary material for "Phase separation in fluids with many interacting components": SI Appendix

#### Supplementary Information

The supplementary information for this manuscript is arranged into separate sections for theory and simulation and includes 5 supplementary figures.

#### Theory

**Thermodynamic model:** The mean-field description of regular solutions is used to study a $N+1$ species fluid mixture, as defined in eq (1), which is reproduced below:

$$f = \sum_{i=1}^{N} \phi_i \log(\phi_i) + \frac{1}{2}\sum_{i=1}^{N}\sum_{j=1}^{N} \chi_{ij}\phi_i\phi_j + \phi_s \log(\phi_s) + \sum_{i=1}^{N} \chi_{is}\phi_i\phi_s$$

Here, $\phi_i$ represent the volume-fraction of each species $i$, and $\phi_s = 1 - \sum_i \phi_i$ is the volume fraction of the remaining component. This remaining species can be interpreted as a solvent, or, another species - if the total solute concentration is held fixed across phases. For simplicity, we will hereby refer to this component as solvent. $\chi_{ij}$ and $\chi_{is}$ are pairwise interaction parameters that are typically proportional to the pair-wise interaction energies ($\epsilon$ ) between $i, j$ and $i, s$ respectively. Normally, the interaction parameter $\chi_{ij} = \frac{z}{2k_BT}(\epsilon_{ij} - \frac{1}{2}(\epsilon_{ii} + \epsilon_{jj}))$, where $z$ is number of short-range contacts, and $\epsilon_{ij}$ represents pairwise energy between components $i, j$. Since we assume component do not interact with themselves, $\chi_{ij} \propto \epsilon_{ij}$.

**Thermodynamic stability:** A solution is thermodynamically unstable and spontaneously phase separates when small fluctuations around the initial composition lead to further lowering of free-energy i.e. the hessian of the free-energy, also referred to as the Jacobian, has at-least one negative eigen-value. The Jacobian is:

$$J_{ij} = \frac{\partial^2 f}{\partial\phi_i\partial\phi_j} = \frac{\delta_{ij}}{\phi_i} + \frac{1}{\phi_s} + \chi_{ij} - \chi_{is} - \chi_{js}$$

In the limit of inert-solvent ($\chi_{is} = 0$) and initially equimolar solutes ($\phi_i = \beta/N$), the above expression reduces to eq. (2). Note if the whole solution, including the solvent, is equi-molar, then $\phi_i = \frac{1}{N+1}; \beta = \frac{N}{N+1}$. The stability of this matrix depends on the eigen-spectrum of:

$$J = \frac{N}{\beta} I + \chi_{eff}; \lambda_J = \frac{N}{\beta} + \lambda(\chi_{eff})$$

$$\chi_{eff,ij} = \frac{1}{1-\beta} + \chi_{ij}$$

Since the pairwise interactions are drawn i.i.d from an underlying distribution with finite mean and variance i.e. $\chi_{ij} \sim Dist(\nu, \sigma^2)$, the eigen-values of this matrix can be derived from Random-matrix theory. The eigen-values follow the semi-circle distribution between $\pm 2\sigma\sqrt{N}$ (Figure S1), except for a lone eigen-value, whose value is centered around $N(\frac{1}{1-\beta} + \nu)$ (1).

The limit of stability i.e. marginal stability is determined when the minimum eigen-value $\lambda_{min} = 0$.The corresponding eigen-vector points to the initial direction of evolution for the instability. From the above distribution, we can see there are 2 limits that naturally arise, since the component-entropy contribution is :

1. *Condensation:* If $N\left(\frac{1}{1-\beta} + \nu\right) < -2\sigma\sqrt{N}$, i.e, $\nu < -\frac{2\sigma}{\sqrt{N}} - \frac{1}{1-\beta}$, only then will the smallest eigen-value be determined by the lone-eigen value (Fig S1B). In the limit of equi-molar solution, this requires $\nu \leq -\frac{2\sigma}{\sqrt{N}} - N$, which represents very strong interactions on average. At marginal stability, $\nu_{marg} = -\frac{1}{\beta} - \frac{1}{1-\beta}$. The corresponding eigen-vector, whose vector-angle with respect to a vector of $(1,1,\ldots,1)_N$ is depicted in gold in Fig S1C, is largely parallel or anti-parallel this vector (angles around $0$ or $180$). Hence, the initial instability direction is either parallel or anti-parallel to $(1,1,\ldots,1)_N$ – which points to an instability that either increases or decreases the volume-fraction of *all* components. This behaves very similar to a bulk fluid undergoing phase separation and is named the condensation transition. More intuitively, if the **average** interaction between species is highly attractive, the fluid undergoes phase separation without changing composition since all the species like to remain inter-mixed. Rather, the interacting components phase separate away from the inert solvent. Overall, this condensation instability requires $\nu < \min\left(-\frac{2\sigma}{\sqrt{N}} - \frac{1}{1-\beta}, -\frac{1}{\beta} - \frac{1}{1-\beta}\right)$, which requires $\nu \sim N$ for equimolar solutions, which represent highly attractive interactions. Note that the 2nd smallest eigen-value is often separated by a band-gap, which corresponds to $\Delta = 2\sigma\sqrt{N} - N\left(\frac{1}{1-\beta} + \nu\right)$ (Figure S1B). In this study, we will focus on de-mixing instability, where there are often multiple eigen-modes beyond the marginally stable point.

2. *Demixing:* Contrary to (1), if $\nu > -\frac{2\sigma}{\sqrt{N}} - \frac{1}{1-\beta}$, then the minimum eigen-value will be determined by the edge of the Wigner semi-circle distribution, which is $-2\sigma\sqrt{N}$. Thus, the

point of marginal stability is determined by: $\sigma_{marg} = \frac{\sqrt{N}}{2\beta}$. The corresponding eigen-vector is determined almost surely perpendicular to the lone eigen-value, and thus has roughly equal number of components with opposite signs (Fig S1C, purple distribution of angles ~ 90 to the vector of ones). Thus the direction of instability points to 2 phases that have distinct compositions and referred to as a demixing transition. Note that for demixing to occur, the constraint on standard deviation of interactions is slacker than the condensation transition: $\sigma \sim \sqrt{N}/2\beta$. For values of $\sigma > \sqrt{N}/2\beta$, multiple negative eigen-values likely contribute to the observed phase behavior, as made vivid in Figures 2,3.

While the stability analyses describes whether a solution will spontaneously phase-separate (spinodal decomposition) and characterize the initial modes of instability, we require a description of dynamics that captures the spatio-temporal evolution of the system, which we describe in detail in the subsequent subsection below.

**Counting the number of unstable modes:** In general, for solutions that are beyond the limit of stability, the number of negative values can be found as the probability of finding eigen-values that off-set the entropic cost ($N/\beta$) multiplied by the total number of eigen-values. This can be written as:

$$< N_{ph} > = N \times F_{2\sigma\sqrt{N}}\left(\lambda \le -\frac{N}{\beta}\right) + 1.$$

Here, $F$ is the cdf of the semi-circle distribution with eigen-values between $\pm 2\sigma\sqrt{N}$, defined as:

$$F_{2\sigma\sqrt{N}}\left(\lambda \le \frac{N}{\beta}\right) = \left(\frac{1}{2} + \frac{\sqrt{N\left(4\sigma^2 - \frac{N}{\beta}\right)}}{4\beta\pi\sigma^2} - \frac{\sin^{-1}\left(\frac{\sqrt{N}}{\beta 2\sigma}\right)}{\pi}\right); \ \frac{\sqrt{N}}{2\beta} < \sigma$$

A useful approximation is made possible by noting that the eigen-values of the semi-circle distribution are uniformly distributed on average. Thus, the average-spacing between eigen-values (when $\nu = 0$) is $\Delta = \frac{4\sigma\sqrt{N}}{N-1} = \frac{4\sigma\sqrt{N}}{N-1}$ – this is because there are roughly N-1 eigen-values in the semi-circle and one lone eigen-value that scales as $\frac{N}{1-\beta} \approx N^2$ for equimolar solutions. Hence the number of eigen-values of the Jacobian that offset the entropic cost can be approximated as:

$$N_{\lambda_J<0} \approx \frac{2\sigma\sqrt{N} - \frac{N}{\beta}}{\Delta} = \frac{N-1}{2}\left(1 - \frac{\sqrt{N}}{2\sigma\beta}\right)$$

The corresponding number of phases is $N_{ph} \approx \frac{N}{2}\left(1 - \frac{\sqrt{N}}{2\sigma\beta}\right) + 1$, which in the limit of strongly variable interaction approaches a theoretical maximal limit of $\frac{N}{2} + 1$. Note that the linear scaling when $\sigma \propto \sqrt{N}$ holds true even without the approximation, as seen from the cdf.

**Optimal number of phases in ensemble with fixed $\sigma$:** In ensembles where the interaction distribution is fixed ($\sigma$ is fixed; or equivalently, the number of sites L, as defined in Fig S5A is fixed) and the initial solution is equimolar ($\phi_i = \frac{1}{N+1}; \beta = \frac{N}{N+1}$), the number of co-existing phases at steady-state follows as:

$$N_{ph} \approx \frac{N}{2}\left(1 - \frac{\sqrt{N}}{2\sigma\beta}\right) = \frac{N}{2}\left(1 - \frac{N+1}{2\sigma\sqrt{N}}\right)$$

To identify the optimal number of phases with changing number of components, we differentiate w.r.t. $N$ and set the derivative to 0:

$$\frac{\partial N_{ph}}{\partial N} = 0 = 3X^2 - aX + 1 = 0$$

Where $X = \sqrt{N}$ and $a = 2\sigma$. Solving these equations show that the one-root is less than $1$, which is not feasible, and the other root $N_{components}^{opt} = \frac{16}{9}\sigma^2\left(1 - \frac{3}{8\sigma^2}\right) \approx \frac{16}{9}\sigma^2$ is the feasible solution that corresponds to an maxima.

**Dynamic model of phase-separation:** The spatio-temporal evolution of the $N$ independent volume-fractions is written following Model B dynamics with diffusive fluxes proportional to the gradient of the chemical potential defined as:

$$\mu_i^b = \frac{df}{d\phi_i} = \log(\phi_i) - \log(\phi_s) + \sum_j \chi_{ij}\phi_j$$

$$flux_i = -M_i \nabla \mu_i^b$$

$$d_t\phi_i + \nabla . flux_i = 0$$

The complete set of dynamical equations are then solved numerically using techniques discussed in the next section. The choice of mobility parameter is decided to be $M\phi_i$, since in the limit of dilute, non-interacting solute ($\phi_i \ll 1, \chi_{ij} = 0, \phi_s \approx 1 - \phi_i$), the flux reduces to Fickian diffusion:

$$flux_i \approx -M\phi_i\left(\frac{1}{\phi_i}\nabla\phi_i - \frac{\nabla\phi_s}{1-\phi_i}\right) \approx -M\nabla\phi_i + O(\phi_i)$$

A similar result holds approximately when $M_i = M\phi_i(1-\phi_i)$, which we find gives results that agree with $M_i = M\phi_i$ (Figure S3B). Finally, the interfacial stabilization terms are added to the chemical potential as a component-independent gradient penalty term ($\kappa > 0$):

$$\mu_i = \log(\phi_i) - \log(\phi_s) + \sum_j \chi_{ij}\phi_j - \frac{\kappa}{2}\nabla^2\phi$$

These above terms can be modified to include reactions as well, an example of which, is provided in the subsequent section.

**Stability analyses:** The stability analyses can be performed in general for the reaction + phase-separation equations as follows (note we will use subscripts of $j$ so as not to confuse with the imaginary square root of $-1$ i.e. $i$)

$$\frac{d\phi_j}{dt} = \vec{\nabla}.\left(M_j\,\vec{\nabla}\mu_j\right) + \sum_k r_{jk}$$

By linearizing around the initial uniform volume-fractions $\phi_j(\vec{r}) = \phi^0 = \beta/N$, we can track the evolution of small fluctuations, written in the form of its inverse fourier-transform $\hat{j} = \phi^0 + \sum_q \mathrm{A_j^q}\exp(-i\,\vec{q}.\vec{r})$. Employing the orthogonality of fourier modes, assuming equimolar solutions without reaction, and linearizing the equation gives the evolution of the amplitude of the $q^{th}$ mode for the $j^{th}$ component evolves as:

$$d_t A_j^q(t) = -\frac{Mq^2\beta}{N}\sum_k J_{jk}A_j^q - \frac{Mq^4\beta}{N}\kappa A_j^q$$

The coupled set of modes, across components, evolve as:

$$d_t\ \overrightarrow{A^q} = -\frac{Mq^2\beta}{N}(J + q^2\kappa\, I)\overrightarrow{A^q}$$

The solution of this linearized equation is $\overrightarrow{A^q}(t) \propto \sum \exp\,(-\frac{Mq^2\beta}{N}\lambda_q(J + q^2\kappa I)t)$ These set of equations are stable across all modes only if all the eigen-values of $J$ are positive since $\kappa > 0$. If the system undergoes phase separation i.e. $\lambda_{min}(J) < 0$, then for all short-wavelength (long length-scales) modes s.t. $q < -\lambda_{min}/\kappa$, the instability propagates, but for larger wavelengths (smaller length-scales), the interface is stabilized. The typical time for an instability to propagate macroscopically ($q \sim 2\pi$ is the shortest wavelength mode under simulation parameters) $t_c \approx \frac{N}{4\pi^2 M\beta\lambda_{min}(J)}$. When active turnover is present, as in eq. (4), the coupled modes evolve subject to:

$$d_t\ \overrightarrow{A^q} = -(k_{off} + \frac{Mq^2\beta}{N}(J + q^2\kappa\, I))\overrightarrow{A^q}$$

Here, we see that the active turnover introduces wave-length independent stabilization of the above equations. More specifically, in the limit of small interface, then the solution becomes asymptotically stable when shortest wavelength modes $(q \sim 2\pi)$ decay i.e. $k \gg \frac{M\beta}{N|\lambda_{min}(J)|}$. More generally, when $k$ is finite, it stabilizes only *certain* eigen-modes at short-wavelengths such that:

$$\lambda \forall |\lambda| < \frac{k_{off}N}{M\beta 4\pi^2}$$

While those eigen modes are stabilized, the remaining modes whose magnitude are larger and directions are largely orthongonal to each other continue to grow, leading to phase separation. Since eigen-modes of the jacobian are equi-spaced on general, this relation can be transformed to give an approximate relation between rate of degradation and number of co-existing phases:

$$n_{phases,k} - n^{ss}_{phases,k=0} = -\frac{k\sqrt{N}}{16\pi^2\sigma}$$

Surprisingly, this linearized relation derived with the assumptions outlined above matches quite well with simulation results (Figure 4B).

#### Simulations

**Numerical implementation:** The set of dynamical equations formulated in Eq. (3) are solved to follow the spatio-temporal evolution of the volume-fractions. All simulations are performed on a 2-D grid of size $L \times L; L = 64$, with a typical time-step $dt = 5e - 6$. The initial conditions are the specified equi-molar volume-fractions with uniform noise in composition that are uncorrelated across spatial positions, with a magnitude typically $\frac{1}{10}$ the mean volume-fraction. Following (2, 3), we implement a implicit (linear or L) -explicit (non-linear or NL terms) formulation to solve for the discrete-time evolution of the volume fraction-fields as follow:

$$\frac{\phi_i(t+\delta t) - \phi_i(t)}{\delta t} = NL_i(\{\phi_i(t)\}) + L_i\big(\phi_i(t+\delta t)\big)$$

$$NL_i(\{\phi_i(t)\}) = \nabla.\big(M\phi_i \nabla \mu_i(\{\phi_i\})\big) + AM\nabla^4\phi_i$$

$$L_i(\phi_i) = -AM\nabla^4\phi_i$$

Here, the fourth-order term is added to ensure numerical stability. The value of $A$ is empirically chosen as $\approx 0.01\max(\chi_{ij})$. All derivative terms and fluxes are computed in fourier space, all other operations are computed in real-space, and fast fourier transforms are used to go between representations. The IM-EX formulation can be solved in fourier form to yield:

$$\hat{\phi}_i(t+\delta t) = \frac{\hat{\phi}_i(t) + \hat{N}_i(\phi_i(t))\delta t}{1 + AMq^4\delta t}$$

Here the fourier transform is defined as: $\hat{f} = \frac{1}{V}\int_V f(\vec{r})\exp(-i\,\vec{q}.\,\vec{r}\,)dr$ . All simulations are run with $\delta t = 5e-6, \kappa = 0.01, M = 1.0$ and run for $2 \times 10^6$ steps to ensure convergence to steady-state, except for the case below. To highlight convergence to steady-state, we run longer simulations (reported in Fig S3C) for $10^7$ steps that highlight convergence by $\sim 10^6$ steps. In typical simulations, $\chi$ is sampled randomly from a normal distribution of given mean, variance and made symmetric by replacing the lower diagonal values with their upper-diagonal counterparts. Simulations are also run with expanded grids to confirm that the observed results are mesh-independent (Fig S3A).

**Analyses of simulation data:** The main observables reported in this study are the number of co-existing phases, their compositions, and the dynamical properties. A given distribution of volume-fractions over the mesh, of the form $N \times L \times L$, contains the information about co-existing phases, where each phase is characterized by its composition (Figure 1B, Figure 2B). The bulk compositions are effectively $k$ significant attractors/vertices in the $N$ component composition-space (Figure 2B). The interfacial regimes represent connecting lines, which in general are curves rather than straight lines, between the various vertices. To count the distinct phases, we first filter out regions that are interfaces (characterized by $\max(\nabla\phi_i)^2 > thresh$ ) and flatten the remaining points into a matrix $X; N \times r$, where r are the number of mesh-points in different bulk-phases.

Subsequently, PCA is performed on $X$ and the $k$ strongly non-zero eigen-modes, which correspond to individual phases, are counted (Figure 1B). Ideally, each mode that is not a phase would have an eigen-value that is zero and vice-versa. In practice, we count the number of eigen-values of $X$ that are greater than $\approx 1e-2$. The logic for this threshold is that if the underlying compositions are randomly sampled around the mean with a typical variance of order of homogeneous concentration i.e. $\sigma \sim 0.1$, eigen-value below $\sigma^2$ are likely to arise from random considerations alone (Marchenko-Pastur distribution). Thus only, compositions that vary more strongly than the stipulated $\sigma$ correspond to bonafide phases. In practice, this approximation captures multiple phases, validated by theory and visualization of simulation data. Then, every spatial point in $X$, of size $N \times 1$, is classified into 1 of $k$ phases using K-means clustering and the cluster-centers are identified, which correspond to typical compositions ( $< \vec{\phi} >_k; N \times 1$) for each of the k-phases. With this, the reported data is obtained by

:

1. *Phase labels:* In plots where phase labels are plotted, each point on the mesh is assigned a phase-label from $1, \ldots, k$ depending on which phase's typical steady-state composition it is closest to ($< \vec{\phi} >_{k,ss}$; as computed by the Euclidean distance). For Figure 2E, the bulk compositions of phases are computed at steady-state, and the labels are assigned at different time-points based on the proximity to steady-state phase compositions. In addition, points that are close to the initial composition $\phi_i = \beta/N$ do not belong to one of the $k$ phases.

2. *Component-enrichment:* For each phase $\gamma$, the number of enriched components is defined as sum of those species whose partition-ratio $p_i^\gamma = \frac{(<\phi_i>_\gamma)}{\phi_i^0} > 1 + \epsilon$, where a small value of $\epsilon = 0.2$ is used to weed out components that are roughly close to their bulk volume-fractions in the initial mixture and the particular phase. The probability distribution shown in Figure 2D and Figure S3 ($p(N_{enriched})$) is derived from data collapsed from multiple co-existing phases per trajectory and across 50 replicate trajectories which only differ in interaction matrix, which is initialized randomly for each replicate.

3. *Compositional similarity:* For any two phases $\gamma, \delta$ the degree of compositional similarity is computed as the angle between the vectors defined by their steady-state compositions $< \vec{\phi} >_{\gamma,ss}, < \vec{\phi} >_{\delta,ss}$. The probability distribution shown in Figure 2C and Figure S3 ($p(\theta)$) is derived from data collapsed from all pairs of co-existing phases per trajectory and across 50 replicate trajectories which only differ in interaction matrix, which is initialized randomly for each replicate.

**Supplementary Figure Captions**

**Figure S1: Eigen-spectrum of jacobian matrix dictates initial modes of instability at spinodal**

A. The eigen-spectrum of a equimolar solution undergoing demixing type instability (N=20, $\chi \sim D(\nu = 0, \sigma = 4.8)$, normalized by $\sigma\sqrt{N}$ and shifted by the entropic costs. The green distribution are values sampled from multiple independent realizations of the Jacobian matrix and the black curves are theoretical predictions. Minimum eigen-values are $< 0$.
B. The eigen-spectrum of a equimolar solute mixture ($\beta = 0.1$) undergoing condensation-type instability (N=20, $\chi \sim D(\nu = -6, \sigma = 1.2)$, normalized by $\sigma\sqrt{N}$ and shifted by the entropic costs. The green distribution are values sampled from multiple independent realizations of the Jacobian matrix and the black curves are theoretical predictions. Minimum eigen-values are $< 0$.
C. The direction of the initial instability is characterized by the angle between the eigen-vector of the smallest eigen-value of the Jacobian and a vector of $(1,1,1 \ldots.,1)_N$. Demixing instabilities are largely orthogonal (purple, centered around 90) while condensation instabilities are either parallel or antiparallel (gold, centered around 0 or 180) to the initial composition i.e. splits into two phases of largely different or similar compositions. The individual points constituting the histogram come from single realizations of the Jacobian.

**Figure S2: Random mixtures exhibit multi-phase co-existence**

Left panel depict volume-fraction profiles of 12 components (labeled $c0$ to $c15$) at steady-state from a single trajectory with identical color-bar scales with an initially equimolar solution with $N = 12$, $\chi \sim Norm(\nu = 0, \sigma = 5.2)$. Darker colors represent regions of higher volume-fraction. Top-right panel depicts the different phases (labeled 1 till 3) present at steady-state. The partition-ratio (ratio of average volume-fraction in a phase over total initial volume-fraction) of all components are plotted for each phase (x-axis) at steady-state.The highlighted components are enriched in those respective phases and the dashed-lines represent no enrichment (partition=1).

**Figure S3: Randomly interacting fluid mixtures show statistical convergence in number of co-existing phases, compositional heterogeneity, and staged phase separation kinetics**

A-C. Simulations tracking number of phases (y-axis) over time (x-axis) for a fixed interaction parameter-set (same as Figure 2) exhibit similar dynamics irrespective of mesh resolution (A), choice of mobility parameter (B), or number of simulation-steps (C). In all cases, solid-lines represents mean of 50 different trajectories, the filled regions represent 1 standard deviation.

D. Number of co-existing phase (y-axis) versus simulation time (x-axis, log-scale) for simulation conditions as in Figure S2. The solid line represents mean of 50 different trajectories, the filled regions represent 1 standard deviation, and the green line represents the specific trajectory whose steady-state properties are shown in Figure S2.
E. Probability (pdf) and cumulative distribution (cdf) of angles between co-existing phases at steady-state for simulation parameters in (D).
F. Probability ($p_{N_{enr}}$) distribution of number of enriched components (x-axis, $N_{enr}$) per phase at steady-state for simulation parameters in (D).

**Figure S4: Stability analyses reveals that phases take characteristic times to emerge macroscopically**

A. Probability (pdf) and cumulative distribution (cdf) of angles between eigen-vectors of the Jacobian matrix across different realizations.
B. Ratio of median time for the macroscopic formation of the $k^{th}$ phase (x-axis,k=1,2,..6), derived from replicate trajectories for same conditions reported in Figure 2, to ratio of predicted time for macroscopic emergence (SI Appendix). The dashed line represents a value of 1 and the error bar represents the standard-deviation in simulation data.

**Figure S5: A simple scaling predicts equilibrium number of phases across diverse parameter regimes**

A. Schematic depicts biophysical model of interactions between species. Each component has L sites of type A or B, with A sites being present at probability $p$. Unlike sites interact favorably with energy $-k\epsilon$ and like sites have unfavorable interactions $\epsilon$. The scaling of the mean and variance of pair-wise interactions between two components with random distributions of A or B sites when $p = \frac{1}{2}, k = 1$ is approximately normal with zero mean and variance proportional to L. Ensembles are as defined in Figure 3B.
B. Variation of number of co-existing phases at steady-state with total solute volume-fractions ($\beta$) for $N = 16, \sigma = 4.8$. Solid lines represent theoretical predictions, dots represent mean of simulation results, and vertical dashes represent one standard-deviation around mean.

C-F. Theoretical predictions of scaling of number of steady-state phases versus number of components in the $\alpha$ (C) and the constant $\sigma$ (E) ensembles. Darker lines represent higher values of $\alpha$ and $\sigma$ respectively. Solid lines are theoretical predictions of scaling of number of steady-state phases versus $\alpha$ (D) and $\sigma$ (F). Darker lines represent higher values of $N$ and dashed-lines represents the predicted upper bound of $\frac{N+1}{2}$ co-existing phases.

**Figure S1: Eigen-spectrum of jacobian matrix dictates initial modes of instability at spinodal**

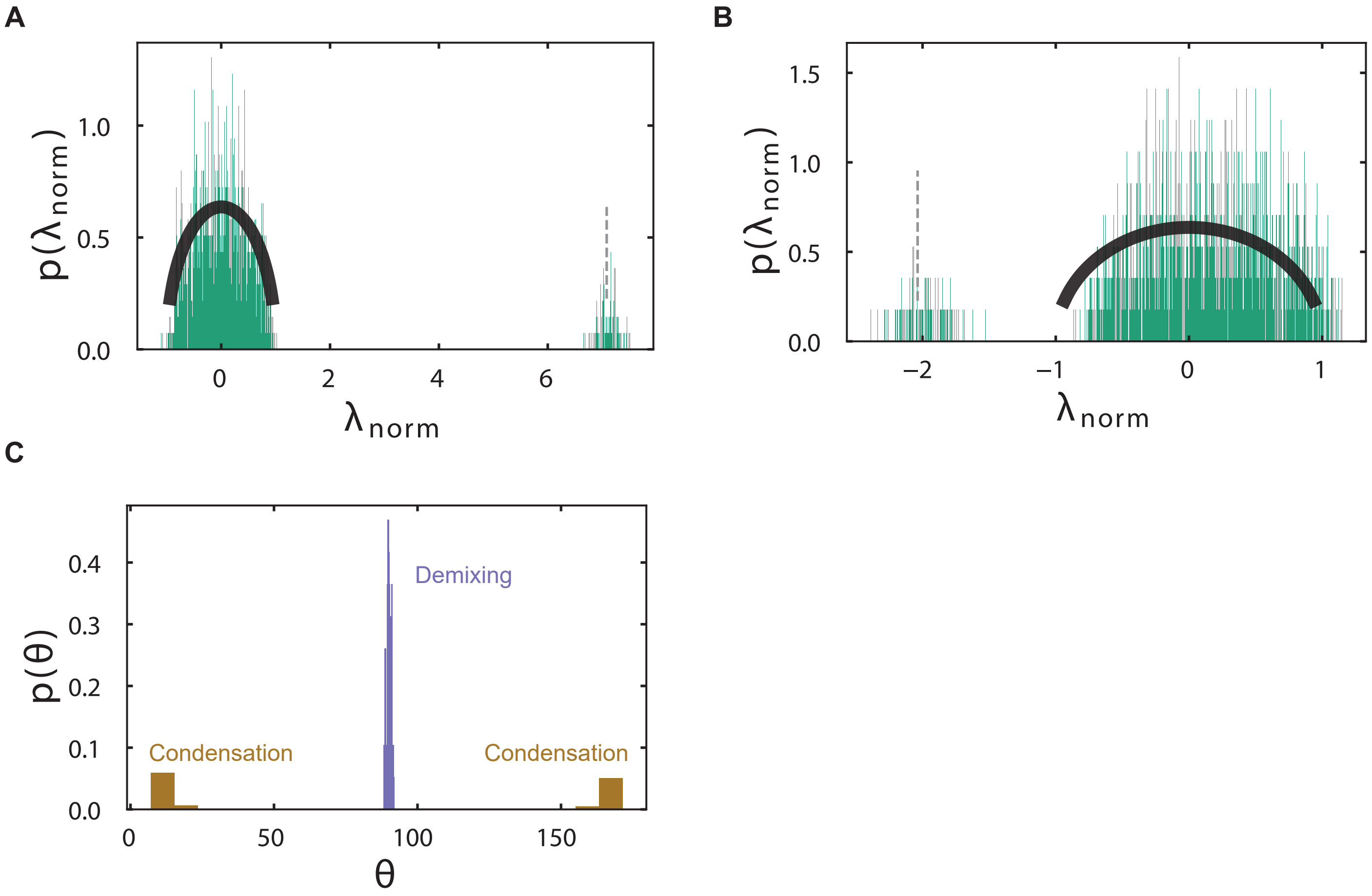


**Figure S1: Eigen-spectrum of jacobian matrix dictates initial modes of instability at spinodal**

**A.** The eigen-spectrum of a equimolar solution undergoing demixing type instability (N=20, χ~Norm(ν=0,σ=4.8)), normalized by σ√N and shifted by the entropic costs. The green distribution are values sampled from multiple independent realizations of the Jacobian matrix and the black curves are theoretical predictions. Minimum eigen-values are <0 indicating presence of instability.

**B.** The eigen-spectrum of a equimolar solute mixture (β=0.1) undergoing condensation-type instability (N=20, χ~Norm(ν=-6,σ=1.2)), normalized by σ√N and shifted by the entropic costs. The green distribution are values sampled from multiple independent realizations of the Jacobian matrix and the black curves are theoretical predictions. Minimum eigen-values are <0 indicating instability.

**C.** The direction of the initial instability is characterized by the angle between the eigen-vector of the smallest eigen-value of the Jacobian and a vector of (1,1,1….,1)_N. Demixing instabilities are largely orthogonal (purple, centered around 90) while condensation instabilities are either parallel or antiparallel (gold, centered around 0 or 180) to the initial composition i.e. splits into two phases of largely different or similar compositions. The individual points in the histogram come from single realizations of the Jacobian.

### Figure S2: Random mixtures exhibit multi-phase co-existence

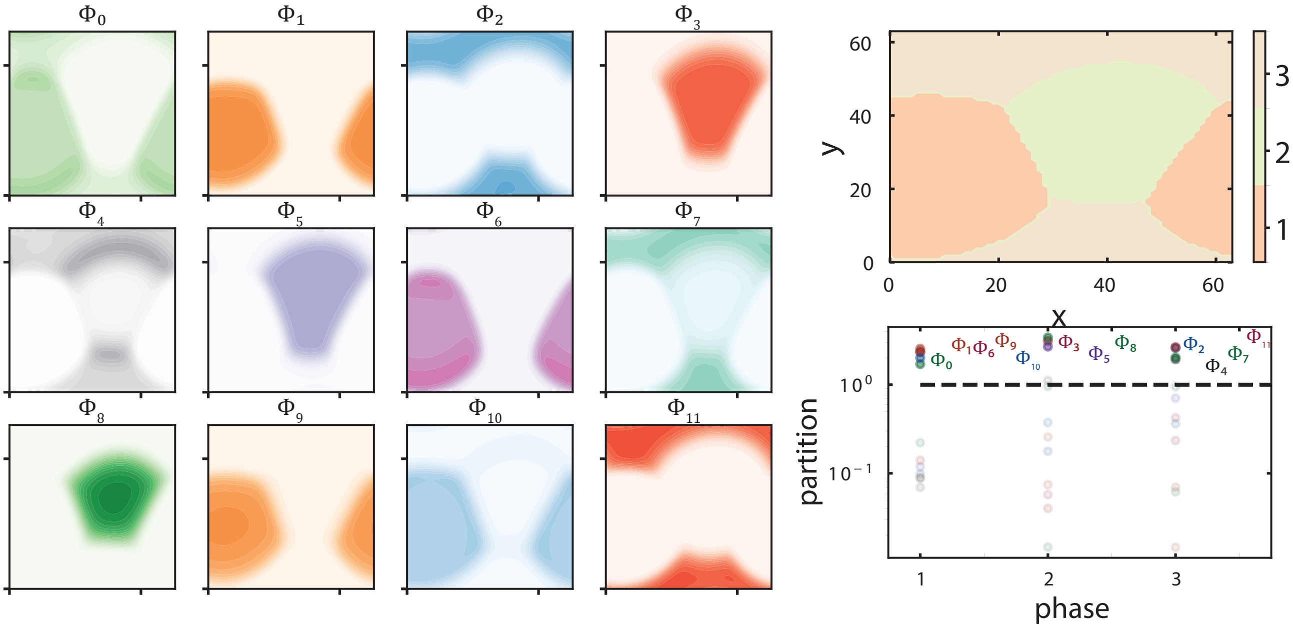


**Figure S2: Random mixtures exhibit multi-phase co-existence**

Left panel depict volume-fraction profiles of 12 components (labeled $\Phi_0$ to $\Phi_{11}$) at steady-state from a single trajectory with identical color-bar scales with an initially equimolar solution with N=12, χ~Norm(ν=0,σ=5.2). Darker colors represent regions of higher volume-fraction. Top-right panel depicts the different phases (labeled 1 till 3) present at steady-state. The partition-ratio (ratio of average volume-fraction in a phase over total initial volume-fraction) of all components are plotted for each phase (x-axis) at steady-state.The highlighted components are enriched in those respective phases and the dashed-lines represent no enrichment (partition=1).

**Figure S3: Randomly interacting fluid mixtures show statistical convergence in number of co-existing phases, compositional heterogeneity, and staged phase separation kinetics**

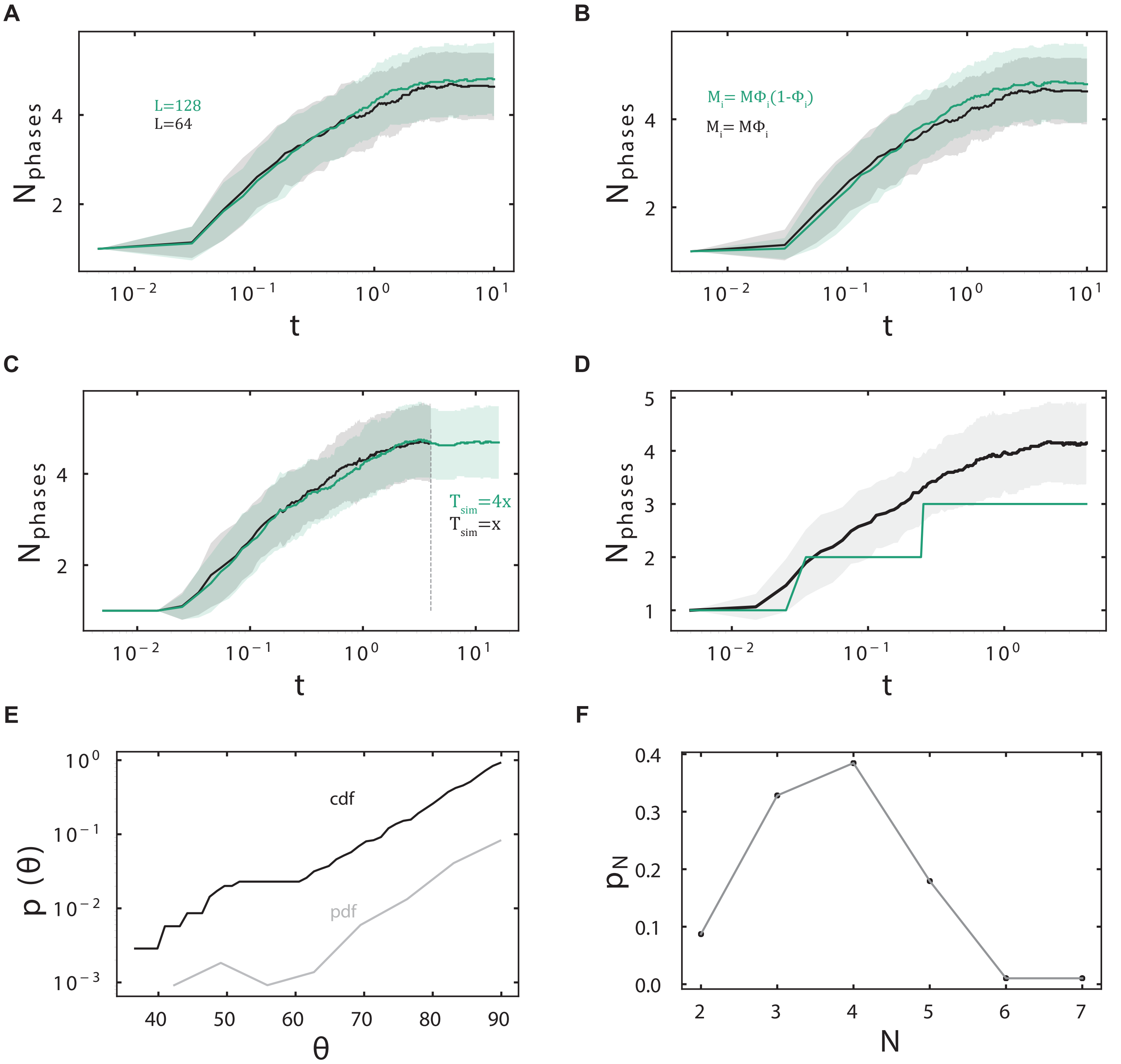


**Figure S3: Randomly interacting fluid mixtures show statistical convergence in number of co-existing phases, compositional heterogeneity, and staged phase separation kinetics**

**A-C.** Simulations tracking number of phases (y-axis) over time (x-axis) for a fixed interaction parameter-set (same as Figure 2) exhibit similar dynamics irrespective of mesh resolution (A), choice of mobility parameter (B), or number of simulation-steps (C). In all cases, solid-lines represents mean of 50 different trajectories, the filled regions represent 1 standard deviation.

**D.** Number of co-existing phase (y-axis) versus simulation time (x-axis, log-scale) for simulation conditions as in Figure S2. The solid line represents mean of 50 different trajectories, the filled regions represent 1 standard deviation, and the green line represents the specific trajectory whose steady-state properties are shown in Figure S2.

**E-F.** Probability (pdf) and cumulative distribution (cdf) of angles between co-existing phases at steady-state (E) and number of enriched components (F) for simulation parameters in (D).

**Figure S4: Stability analyses reveals that phases take characteristic times to emerge macroscopically**

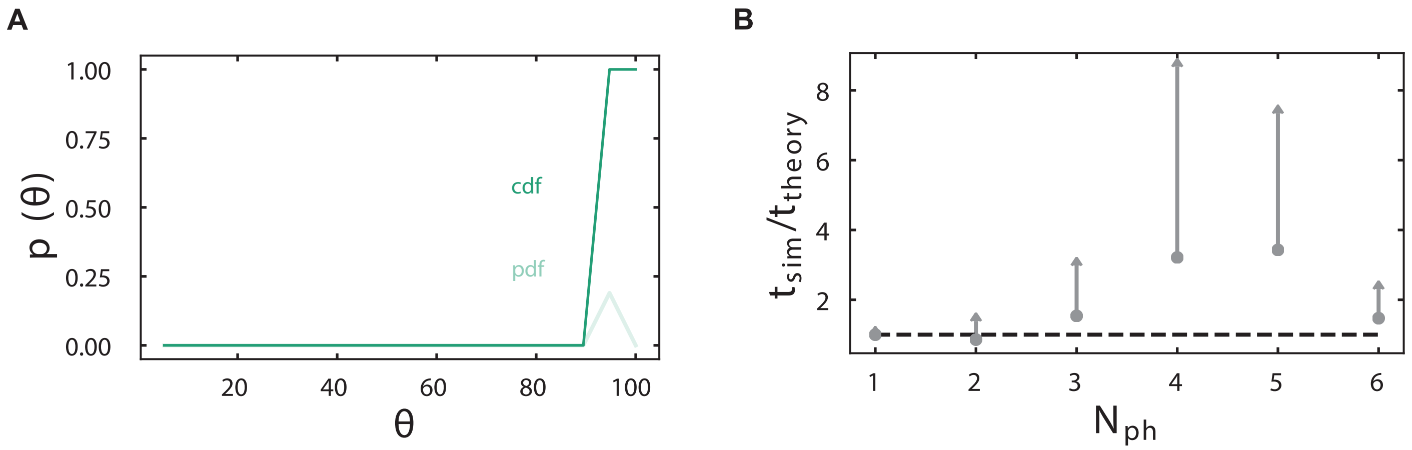


**Figure S4: Stability analyses reveals that phases take characteristic times to emerge macroscopically**

**A.** Probability (pdf) and cumulative distribution (cdf) of angles between eigen-vectors of the Jacobian matrix across different realizations.

**B.** Ratio of median time for the macroscopic formation of the k^th phase (x-axis,k=1,2,..6), derived from replicate trajectories for same conditions reported in Figure 2, to ratio of predicted time for macroscopic emergence (SI Appendix). The dashed line represents a value of 1 and the error bar represents the standard-deviation in simulation data.

**Figure S5: A simple scaling predicts equilibrium number of phases across diverse parameter regimes**

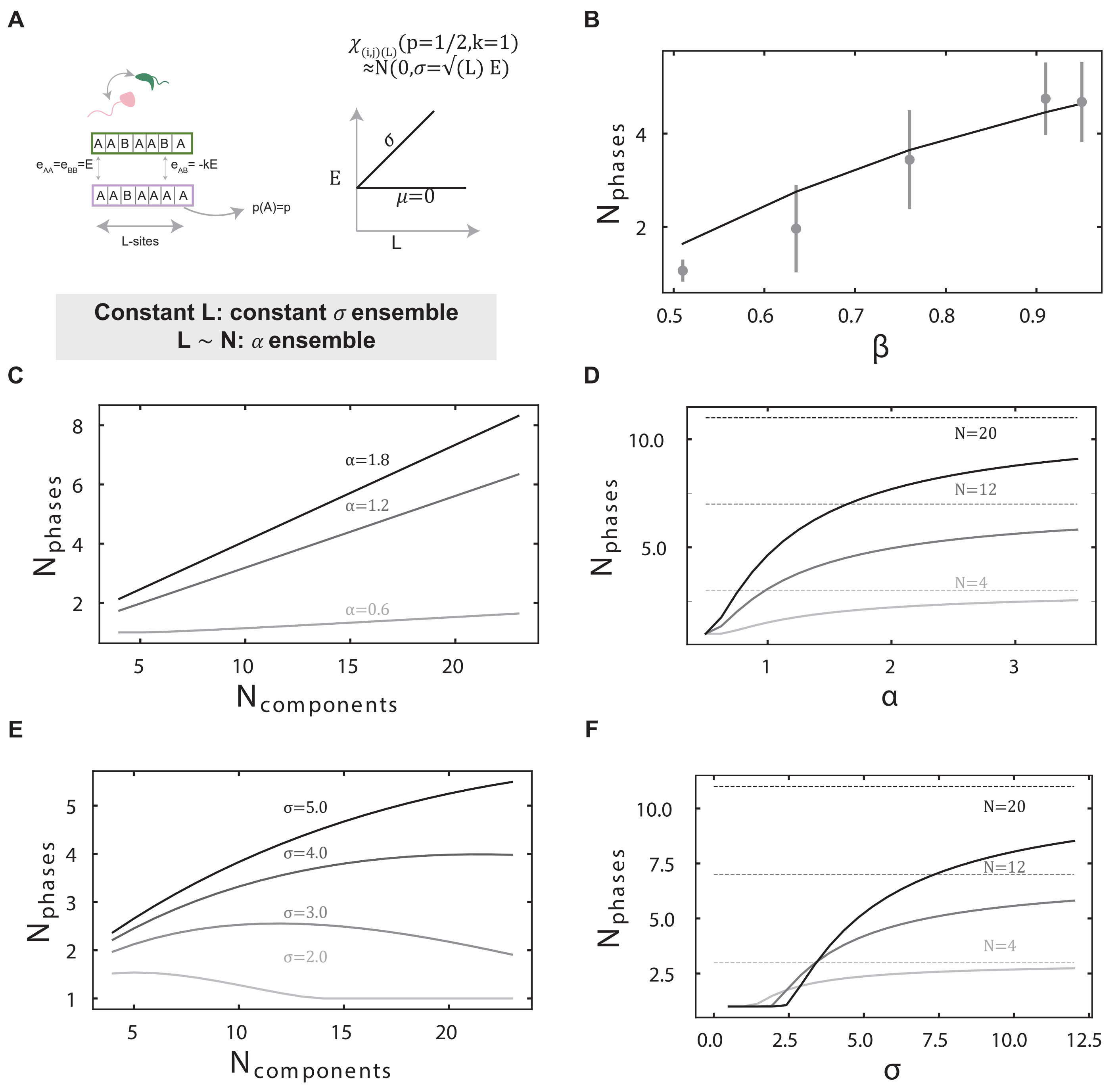


**Figure S5: A simple scaling predicts equilibrium number of phases across diverse parameter regimes**

**A.** Schematic depicts biophysical model of interactions between species. Each component has L sites of type A or B, with A sites being present at probability p. Unlike sites interact favorably with energy -kϵ and like sites have unfavorable interactions ϵ. The scaling of the mean and variance of pair-wise interactions between two components with random distributions of A or B sites when p=1/2,k=1 is approximately normal with zero mean and variance proportional to L. Ensembles are as defined in Figure 3B.

**B.** Variation of number of co-existing phases at steady-state with total solute volume-fractions (β) for N=16,σ=4.8. Solid lines represent theoretical predictions, dots are mean of simulation results, and vertical dashes represent one standard-deviation around mean.

**C-F.** Theoretical predictions of scaling of number of steady-state phases versus number of components in the α (C) and the constant σ (E) ensembles. Darker lines represent higher values of α and σ respectively. Solid lines are theoretical predictions of scaling of number of steady-state phases versus α (D) and σ (F). Darker lines represent higher values of N and dashed-lines represents the predicted upper bound of (N+1)/2 co-existing phases.
